# Mapping of CELF1-RNA interactions reveals post-transcriptional control of lens development

**DOI:** 10.64898/2026.01.09.698617

**Authors:** Justine Viet, Matthieu Duot, Agnès Méreau, Yann Audic, Iwan Jan, David Reboutier, Catherine Le Goff-Gaillard, Sarah Y Coomson, Salil A Lachke, Carole Gautier-Courteille, Luc Paillard

## Abstract

Precise post-transcriptional regulation of gene expression is essential for vertebrate lens development. Disruption of the gene encoding the RNA-binding protein CELF1 leads to early-onset cataract in mice. Here, using iCLIP-seq in lenses, we mapped transcriptome-wide CELF1 binding sites, revealing interactions with the 3′UTRs of key transcripts involved in lens development and pathology like *Gja8*, *Jag1*, *Maf*, *Pax6*, or *Prox1*. Integrated analysis with transcriptomic data and luciferase reporter assays demonstrated that binding of CELF1 protein represses its target mRNAs by destabilizing transcripts and/or inhibiting their translation. Indeed, the cataract-linked genes *Maf* and *Gja8* are upregulated in *Celf1*^cKO^ lenses. In *Xenopus laevis*, overexpression of *maf* resulted in abnormal lens structure and eye morphology, confirming the developmental relevance of CELF1-mediated repression. Our findings uncover a post-transcriptional network in which CELF1 controls lens morphogenesis by limiting the expression of critical genes at the mRNA level to achive their proper dosage.

## INTRODUCTION

The ocular lens is a biconvex organ located behind the iris and in front of the retina. Along with the cornea, the lens focuses light onto the retina, a function that depends critically on its transparency and precisely controlled refractive index. The lens is composed mainly of highly differentiated, non-proliferative lens fiber cells arranged concentrically. Fibers are surrounded anteriorly by a monolayer of epithelial cells. These epithelial cells divide in the proliferation zone and, when they reach the equatorial region of the lens, they initiate differentiation into fiber cells. The specific optical properties of the lens are achieved through multiple mechanisms, including the elimination of organelles and nuclei in fiber cells and the accumulation of highly refractive proteins known as crystallins [1–4].

Loss of lens transparency is defined cataract, a major age-related eye disease and a leading cause of blindness worldwide [5–7]. Cataracts result from the formation of light-scattering protein aggregates in the lens. They essentially affect older individuals as such aggregates gradually accumulate with age. Environmental conditions such as UV light and oxidative stress promote this process [8]. However, cataracts can also be congenital, typically with a genetic basis [7,9]. The Cat-Map database catalogs cataract-associated genes and loci in vertebrates [10,11], listing over 500 genes as of its February 2025 release. These include genes that encode crystallin proteins, membrane proteins such as connexins and the aquaporin MIP, or transcription factors essential for lens development such as *PAX6*, *MAF*, *FOXE3* or *HSF4* [11].

Beyond transcriptional regulators, post-transcriptional mechanisms play a pivotal role in mediating spatiotemporal gene expression control during lens development. RNA-binding proteins (RBPs), in particular, have emerged as critical regulators [12,13]. For example, *TDRD7* was first implicated in cataract through human balanced chromosomal rearrangements and inherited mutations, and validated in animal models [14]. A large genome-wide association study linked several RBP genes, including *THOC7*, *QKI*, *CAPRIN2*, *RBFOX1*, to cataract [15]. Functional studies have shown that conditional inactivation of *Caprin2* or *Qki* in the mouse lens leads to cataract, the latter due in part to disrupted cholesterol biosynthesis [16–18]. Interestingly, although QKI is typically characterized as an RBP [19], it may function at the transcriptional level in the lens [18]. Recent transcriptome/proteome analyses have expanded the list of RBPs expressed in the lens [20].

CELF1 (CUGBP, Elav-like family member 1) is a well-known RBP recognized for its role in alternative splicing regulation [21–25]. It also promotes transcript destabilization via binding to the 3’ untranslated regions (3’UTRs), targeting genes such as *JUN*, *JUNB* or *TNFRS1B* [26–29]. It also controls mRNA translation of mouse *Cyp19a1* as well as of several human genes involved in epithelial-to-mesenchymal transition [30,31]. Conditional deletion of *Celf1* gene in the mouse lens causes early-onset cataracts [32]. In human, CELF1 is indirectly implicated in cataract through Myotonic Dystrophy, type 1 (DM1), a multisystem disorder that causes cataracts among other symptoms. The genetic basis of DM1 is a CTG repeat expansion in the *DMPK* gene, which leads to increased CELF1 activity and reduced MBNL1 function by still incompletely understood mechanisms. CELF1 and MBLN1 both being RBPs involved in splicing control, this would disrupt splicing regulation and cause the symptoms of DM1 [21–23]. A related disease, Myotonic Dystrophy, type 2 (DM2), also presents with cataracts and is caused by repeat expansion in the *CNBP* gene, which encodes a nucleic acid (including RNA) binding protein [33,34].

In the lens, a candidate gene approach revealed that CELF1 regulates the two key transcription factors PAX6 and PROX1 at a post-transcriptionnal level [35]. A recent transcriptomic study further showed that *Celf1* inactivation disrupts lens differentiation, as revealed by the down-regulation of multiple lens terminal differentiation genes [36]. However, the direct molecular mechanisms by which the lack of CELF1 results in cataracts remain unclear. Specifically, the full repertoire of RNAs bound by CELF1 in the lens and the functional consequences of these protein-RNA interactions are unknown.

To address this gap, we performed here transcriptome-wide identification of CELF1-bound RNAs in mouse lenses using iCLIP-seq, a method that maps RNA-protein interactions at nucleotide resolution [37]. By performing an integrated analysis of iCLIP-seq data with gene expression profiling data on *Celf1*^cKO^ lenses, and mechanistically testing key candidates with luciferase reporter assays, we show that CELF1 functions as a global post-transcriptional repressor in the lens. Using a stepwise prioritization pipeline, we identify high-confidence CELF1 ligands whose increased expression in the absence of CELF1 likely contributes to cataract. Among them, we validate *Maf* and *Gja8* as key CELF1 ligands, showing that they are upregulated in the absence of CELF1 *in vivo*. Overexpression of *Maf* in Xenopus larvae elicites lens defects reminicent of those observed in *Celf1*-deficient mice, indicating that tight regulation of *Maf* by CELF1 is important for normal lens development. Together, these findings (i) reveal for the first time the *in vivo* ligands of CELF1 in a specific organ, (ii) demonstrate CELF1’s global repressive role in lens gene expression, and (iii) identify candidate genes whose increased expression is likely pathogenic, offering a complementary resource to traditional loss-of-function screens for cataract gene discovery.

## RESULTS

### iCLIPseq identifies mRNA ligands of CELF1 in mouse lenses

To identify the RNA ligands of CELF1 at the transcriptome-wide level, we performed two independent iCLIP-seq experiments on mouse lenses as described [38]. Briefly, lenses freshly dissected from 2-month-old mice were UV-irradiated to induce covalent RNA-protein crosslink, lysed, gently treated with RNase, and subjected to immunoprecipitation with an anti-CELF1 antibody. The co-immunoprecipitated RNAs were then recovered, reverse-transcribed, amplified and deep sequenced.

The two iCLIP-seq datasets differed in RNase stringency: CLIP2 used a higher concentration of RNase than CLIP1, resulting in a lower number of uniquely mapped reads (Figure 1A). In iCLIP, reads terminate at the site where reverse transcriptase is halted by the cross-linked protein, enabling identification of binding sites at nucleotide resolution [38]. In Figure 1A, “cross-link” refer to individual CELF1-RNA interaction sites; the number of reads mapping to a site determines its score. “Peaks” are statistically significant cross-linking sites, identified by comparison to a randomized distribution [39]. Nearby peaks are grouped into clusters, revealing over 66,000 and nearly 10,000 CELF1 binding clusters in CLIP1 and CLIP2, respectively (Figure 1A). More than 50% of these clusters localize to 3’ untranslated regions (3’ UTR) in both datasets (Figure 1B). We focused our analysis on these 3’UTR clusters, which were found in 6,066 (CLIP1) and 2,561 (CLIP2) genes. Only 199 CLIP2 genes are absent in CLIP1, implying that the CLIP2 geneset is largely included within the CLIP1 geneset. The complete gene list (6,265 genes) with iCLIP-seq data is available as Table S1.

**Figure 1.**
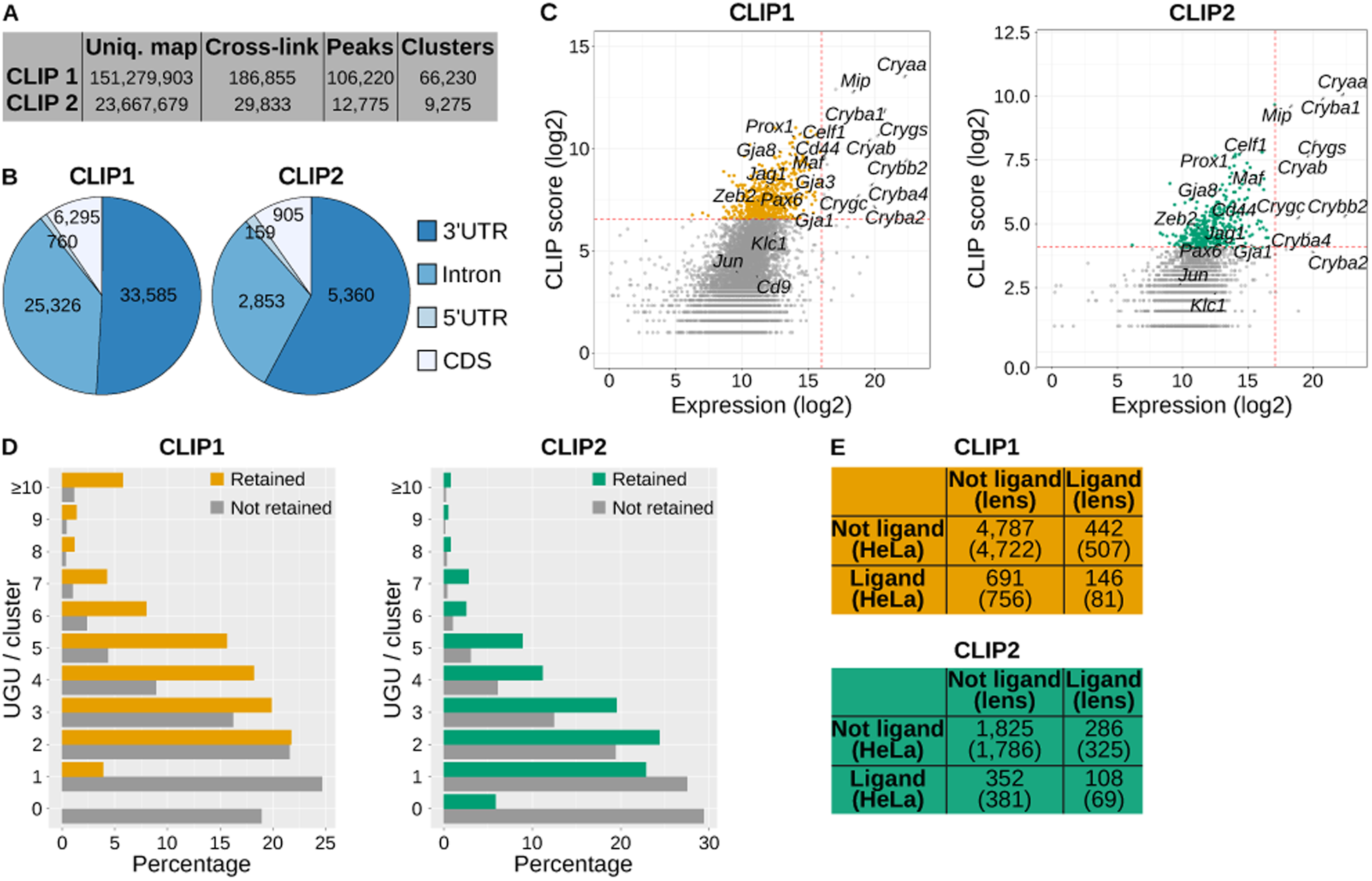
iCLIP-seq of CELF1 in mouse lenses **A**, Summary of iCLIP-seq data showing the number of uniquely mapped reads, cross-link sites, significant cross-link sites (peaks) and peak clusters, as defined in the text, for the two iCLIP-seq experiments. **B**, Pie charts showing the distribution of CELF1 binding clusters across gene regions. Left CLIP1 (33,585 clusters in 3’UTRs of 6,066 genes); right CLIP2 (5,360 clusters in 3’UTRs of 2,561 genes). **C**, CLIP scores plotted against gene expression levels (log scales). Each dot represents one gene. Genes retained as CELF1 ligands based on predefined thresholds (dashed red lines) are shown in orange (CLIP1, N = 588) and green (CLIP2, N = 394). **D**, Distribution of UGU trinucleotide numbers within CELF1 binding cluster. For genes with multiple clusters, only the cluster with the highest UGU count was retained. **E**, Contingency tables comparing CELF1 ligands identified in mouse lenses with those in HeLa cells. CLIP1: 6,066 genes with at least one CELF1 binding cluster in the 3’UTR, of which 588 were retained as ligands. CLIP2: 2,561 genes with at least one cluster in the 3’UTR, of which 394 were retained as ligands. Expected counts under independence are indicated in parentheses. p=5.5e-16 and p=3.6e-08, CLIP1 and CLIP2, respectively, chi-square test.

To quantify CELF1 binding per gene, we defined a “CLIP score” as the sum of scores of CELF1 binding clusters within its 3’UTR. Figure 1C presents a smear plot-like distribution of CLIP scores against gene expression (based on RNA-seq from wild-type lenses [36]). Genes in the top 10% of CLIP scores were retained. We observed that highly expressed genes, particularly crystallins, also showed high CLIP scores, likely due to non-specific binding to very abundant RNAs. To mitigate this bias, the top 0.5% most highly expressed genes were excluded. The retained genes (colored dots in Figure 1C) include 588 (CLIP1) and 394 (CLIP2) genes (Table S1). Notably, several known regulators of lens biology were identified in both datasets, including *Pax6*, *Prox1*, *Gja8*, *Maf*, *Jag1*, or *Celf1* itself. Among previously reported CELF1 ligands [24,25,29,40], *Cd44* was recovered, but *Jun*, *Klc1* and *Cd9* were not detected using our thresholds.

Although CELF1 binding sites are generally poorly conserved, they are enriched in UGU trinucleotide motifs [40]. As a quality control, we compared UGU content in the CELF1 binding clusters of retained versus non-retained genes (Figure 1D). Even non-retained genes are part of the N=6,066 or N=2,561 genes in CLIP1 and CLIP2 that have at least one CELF1 binding cluster in their 3’UTR, albeit considered as non-significant. The CELF1 binding clusters of retained genes show strong UGU enrichment. For example, in CLIP1, 437 retained genes have clusters with at least 3 UGUs, and 151 have 2 UGUs or fewer. Only 360 genes with ≥ 3 UGU were expected by chance, and 227 with ≤ 2 UGU (p=1.2e-77, chi-square test). Similar enrichment was observed in CLIP2 (p=2.4e-21). Hence, the CELF1 binding clusters of retained genes are enriched in UGUs.

To further validate our dataset, we compared CELF1 ligands identified in mouse lens with those reported in human HeLa cells [41]. Of the 588 CLIP1 ligands and 394 CLIP2 ligands, 146 and 108 overlap with CELF1 ligands in HeLa cells, respectively, which is significantly more than expected by chance (Figure 1E, Table S1). Hence, although the repertoires of CELF1 ligands cannot largely overlap between two highly different cell types that express dissimilar sets of genes, the enrichment of previously identified CELF1 ligands within both CLIP1 and CLIP2 support the biological relevance and reproducibility of our findings.

### CELF1 is a global post-transcriptional repressor in the lens

From CLIP1 and CLIP2, we identified 588 and 394 CELF1-bound mRNAs, respectively, with 293 shared between them (Figure 2A). Thus, CLIP2 (high stringency) is largely included within CLIP1 (293 / 394 = 74.6%). We defined two gene sets. The *intersection* includes 293 genes, shared between both CLIP1 and CLIP2, representing high-confidence ligands of CELF1. The *union* includes 689 genes, 293 commonly found in both CLIP1 and CLIP2 plus those exclusively found in CLIP1 (295) and CLIP2 (101) (Figure 2A, Table S1).

**Figure 2.**
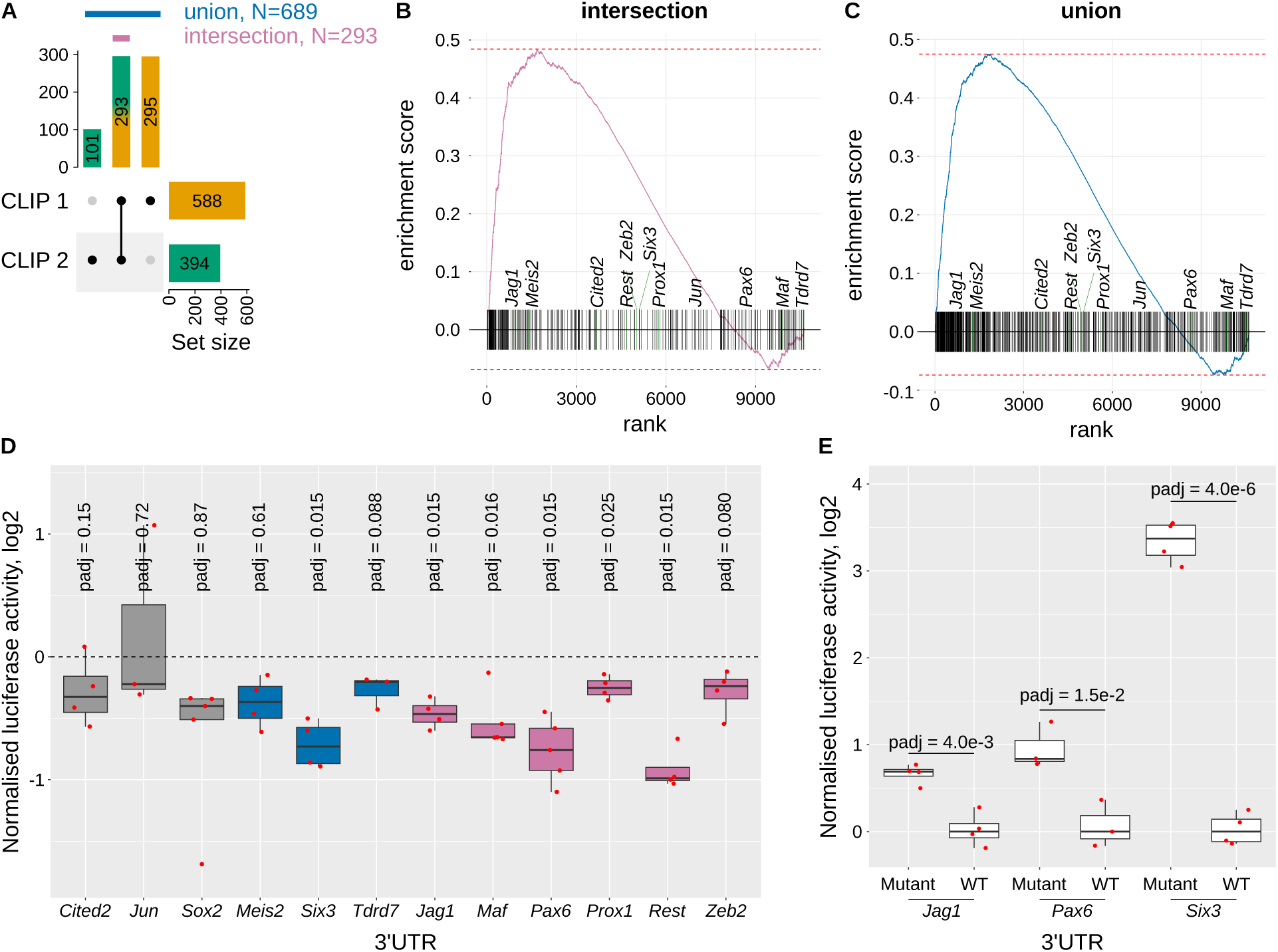
Impact of CELF1 binding on mRNA levels and translation **A**, Upset plot showing the overlap of CELF1 mRNA ligands identified in CLIP1 and CLIP2. The intersection (magenta, N=293) includes mRNAs bound in both experiments, and the union (blue, N = 689) includes mRNAs bound in at least one. **B-C**, GSEA-like plots. Genes were ranked by signed FDR (-log10 scale) for differential expression between wild-type and *Celf1-*CKO lenses [36]. Most significantly up-regulated genes in *Celf1-*CKO lenses are on the left; most significantly down-regulated genes on the right. Vertical lines indicate CELF1 ligands in the intersection (**B**) or the union (**C**). Selected genes used in subsequent experiments are labeled. **D**, Luciferase reporter assays using constructs with the indicated 3′UTRs downstream of firefly luciferase. The same plasmid also encodes a renilla luciferase. Luciferase activity was measured following co-transfection by the luciferase plasmid and a plasmid driving the expression of *Celf1* or *H2B* as a control. Ludiferase was normalized sequentially: (1) to renilla luciferase (F/R), (2) to the F/R ratio of a control (plasmid) 3’UTR, and (3) to the F/R of cells transfected control with H2B. Values significantly below 0 indicate CELF1-dependent repression. *padj*, Benjamini-Hochberg adjusted p-value, Student’s t test (null hypothesis: mean = 0). Grey, 3’UTRs from genes not identified as CELF1 ligands; blue, genes identified as a CELF1 ligand in a single CLIP dataset; magenta, genes identified as a CELF1 ligand in both CLIP experiments. **E**, Luciferase assays comparing wild-type and mutant 3’UTRs. Normalized as in **D**, except without H2B normalization. In **D** and **E**, each dot represents the median of at least 4 technical replicates, with biological replicates shown as separate dots.

We next integrated these datasets with transcriptomic data comparing wild-type and *Celf1*-conditional knock-out (cKO) lenses [36]. Using a GSEA-like approach [42], we observed that the ligands of CELF1 were highly enriched among the genes most significantly up-regulated in the absence of CELF1 (Figure 2B). This was true for both the intersection (ES = 0.48, p = 8.6e-12) and the union (ES = 0.48, p = 2.2e-23), strongly suggesting that CELF1 broadly destabilizes its bound mRNAs in the lens.

We next investigated whether CELF1 affects mRNA translation beyond this effect on mRNA stability. We performed luciferase reporter assays upon *Celf1* overexpression using 3’UTRs from 12 genes (Figure 2D). These included 3’UTRs from (i) 3 genes not identified as CELF1 ligands (*Cited2*, *Jun*, *Sox2*), (ii) 3 genes identified as a CELF1 ligand in a single CLIP dataset (*Meis2*, *Six3*, *Tdrd7*), and (iii) 6 genes identified as a CELF1 ligand in both CLIP experiments (*Jag1*, *Maf*, *Pax6*, *Prox1*, *Rest*, *Zeb2*). In these experiments, luciferase activities significantly below 0 reveal translational repression by the overexpression of *Celf1* [30]. Translation was not repressed by *Celf1* overexpression for category (i) (non-ligand) 3’UTR. It was repressed for only one 3’UTR ( *Six3*) belonging to category (ii). However, it was repressed for 5 out of 6 3’UTRs belonging to category (iii): *Jag1*, *Maf*, *Pax6*, *Prox1*, *Rest*.

To confirm that repression is due to CELF1 binding, we examined the distribution of CELF1 binding clusters along these 3’UTRs. While CELF1 spreads all along the 3’UTR for most of them, we identified discrete high-density regions of CELF1 clusters in *Jag1*, *Pax6* and *Six3* (Figure S1). Deleting these regions from the 3’UTRs relieved at least partly translational repression (Figure 2E), demonstrating that CELF1 represses translation via direct binding.

Interestingly, only *Jag1* and *Meis2* among the 12 tested 3’UTRs are in the GSEA leading edge (Figures 2B-C, most significantly up-regulated expressed genes in the absence of CELF1). However, luciferase assays indicate that CELF1 represses translation of *Six3*, *Maf*, *Pax6*, *Prox1*, or *Rest* (Figure 2D), even though their mRNA levels are not increased in the cKO lenses (Figures 2B-C). Thus, CELF1 likely regulates more targets at the translational level than suggested by RNA-seq alone. Together, these findings highlight CELF1 as a major post-transcriptional repressor in the lens, regulating gene expression through both mRNA stability and/or translation.

### CompBio-based analysis of iCLIPseq-based CELF1 mRNA ligands

We used a computational tool called CompBio (*Comprehensive Multi-omics Platform for Biological InterpretatiOn tool*) to identify biological themes enriched in the iCLIP union ligands (689) [43–47]. CompBio identified themes related to “lens development and maintenance”, “connexins, gap junction proteins and intercellular communication”, and “basement membrane composition and capsule organization” that are related to lens biology, suggesting that Celf1 directly binds to mRNAs related to function in the lens (Figure 3). Among other themes, the “connexins, gap junction proteins and intercellular communication” theme may help explain the cellular vacuole defects observed in *Celf1*^cKO^ lenses [32]. Additionally, CompBio also identified themes such “actin cytoskeleton, cell adhesion” and “PAK signaling and cytoskeletal regulation” which may help explain the cytoskeletal defects observed in the *Celf1*^cKO^ lenses [32]. Interestingly, “RNA binding and post-transcriptional regulation”, “mRNA metabolism and regulation of translation” and “post-transcriptional mRNA decay and regulation of mRNA stability” were also identified as themes, suggesting that CELF1 directly binds to mRNA that are themselves involved in these post-transcriptional processes. Additionally, themes related to “neuronal development” and “neuronal cytoskeleton and axonal transport” were also found to be enriched, among other themes. In conjunction with previously described mechanisms [48,49], this suggests that CELF1 contributes to the repression of neuronal genes in the lens. Together, CompBio analysis shows that CELF1 target mRNAs are involved in different aspects of cellular control.

**Figure 3.**
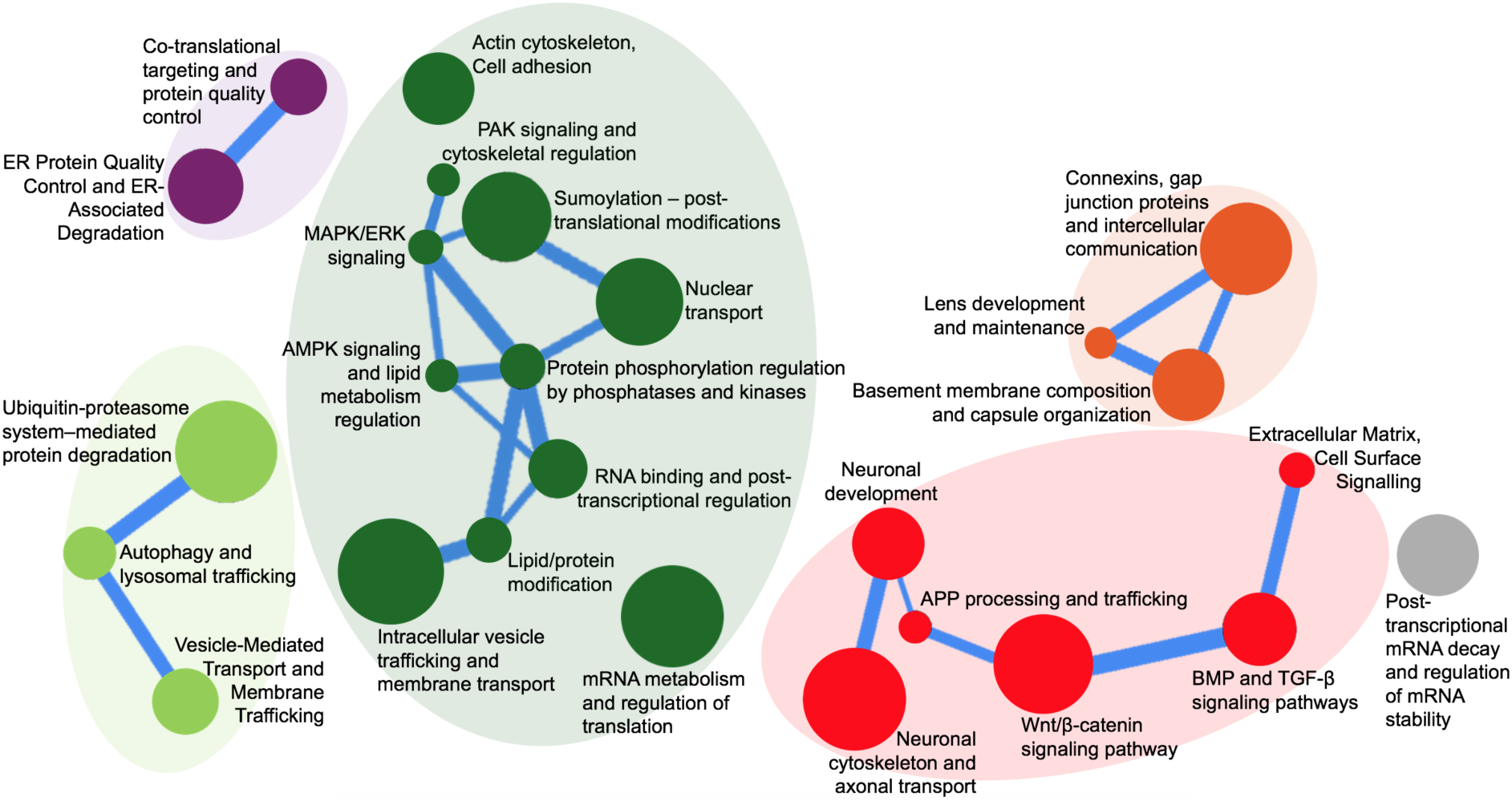
Enriched biological themes identified by CompBio in CELF1 iCLIP datasets in the lens. CompBio-based analysis of 689 genes (found in CLIP1 and/or CLIP2) identifies enriched biological themes. CompBio does de novo analysis of the test list of candidates using the published scientific literature covering >30 million PubMed abstracts and >3 million full-text articles. Circles denote nodes that represent enriched themes while the edges between nodes indicate that genes are shared to denote the different themes they connect. The size of a circle is based on the number of genes that contribute to the node and the thickness of an edge denotes the amount of overlapping genes. In the different clusters, the themes that are related are represented with similar colors.

### CELF1 ligands encode key regulators of lens development

To investigate the functional relevance for lens development and diseases of CELF1 ligands, we first assessed their lens specificity using the iSyTE 2.0 database. The lens-enrichment module of iSyTE 2.0 compares gene expression in the lens and in the whole body, to identify lens-specific and lens-enriched genes [50]. Figures 4A-A’ show higher lens-enrichment for CELF1 ligands than for non-ligands. Hence, CELF1 ligands are enriched in genes that show a preferential expression in the lens, suggesting their functional importance in lens biology.

**Figure 4.**
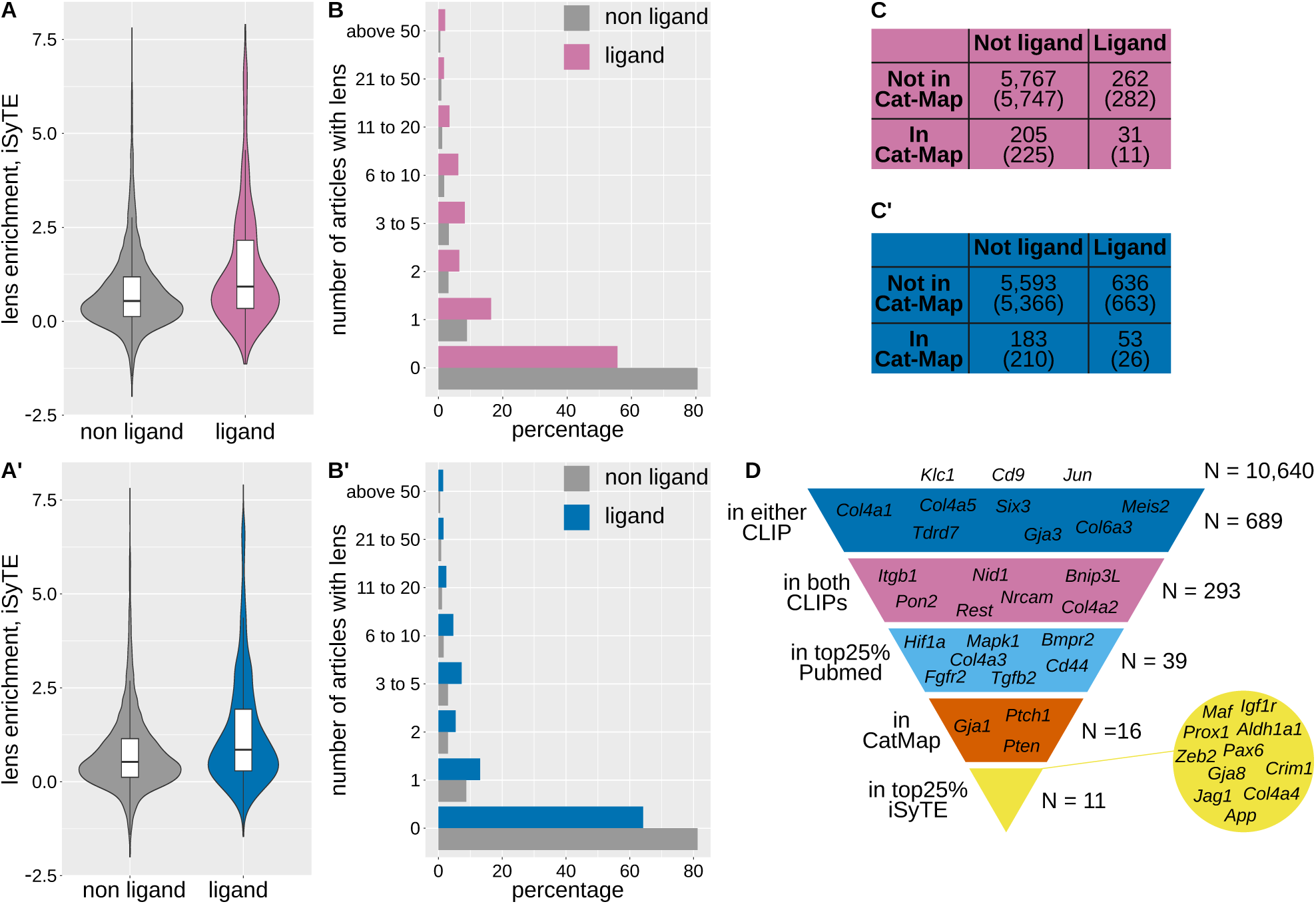
CELF1 ligands are enriched for genes involved in lens development and disease **A-A’**, Lens expression enrichment of CELF1 ligands versus non-ligands, using iSyTE 2.0. **A**, intersection (p = 6.4e-11, Wilcoxon test), **A’** union (p = 1.5e-18). **B-B’**, Number of PubMed citations per gene retrieved with “[gene name] AND lens” for CELF1 ligands and non-ligands. **B**, intersection (p = 1.7e-24, chi-square test comparing the distribution of ligands and non-ligands between no citation and at least one citation), **B’** union (p = 2.0e-25). **C-C’**, Contingency tables comparing CELF1 ligands and non-ligands for inclusion in Cat-Map. **C**, intersection (p = 9.6e-10, chi-square test), **C’** union (p = 4.1e-9, chi-square test). **D**, Gene prioritization workflow (left) and the number of gene passing each selection step (right), with example genes filtered out at each stage.

As an orthogonal approach, we performed PubMed queries combining each gene with the keyword “lens”. With the intersection, we found that fewer ligands than non-ligands do not appear with lens in any article (55.6% of ligands and 80.5% of non-ligand). Ligand genes were more frequently co-mentioned with “lens” than non-ligands, including in multiple publications (Figure 4B). This is also the case when considering the union as the gene ligands (Figure 4B’). Hence, CELF1 ligands are enriched in genes that have already been investigated for their roles in lens biology.

We next examined their association with cataract. According to Cat-Map [10,11], 538 genes are implicated in cataract (February 2025 release). Of these 31 (intersection) and 53 (union) overlap with CELF1 ligands, significantly more than expected by chance (Figures 4C-C’). This implicates CELF1 ligands in lens disease.

### A multistep pipeline prioritizes key CELF1 ligand for functional studies

To prioritize ligands for functional validation, we applied a multistep filtering pipeline (Figure 4D). As previously described, starting from 10,640 genes expressed in the lens, we retained 689 genes that are ligands of CELF1 in at least one iCLIPseq experiment, then 293 genes that are ligands of CELF1 in both. Of them, we retained 39 genes that are in the upper quarter of the number of publications in a PubMed search with “lens”. The next filtration based on the presence in Cat-Map retained 17 genes, including *Gja1* which encodes Connexin 43 required for cell-to-cell diffusion of small molecules in the lens [51], *Ptch1* which encodes a sonic hedgehog receptor involved in radiation-induced cataract at least in animal models [52,53], or *Pten*, which encodes a lipid phosphatase involved in lens ion transport [54]. Finally, the last filtration relying on a high lens-enrichment of expression (top quarter) retained 11 genes: *Aldh1a1*, *App*, *Col4a4*, *Crim1*, *Gja8*, *Igf1r*, *Jag1*, *Maf*, *Pax6*, *Prox1*, *Zeb2* (Figure 4D).

### *Maf* and *Gja8* expression levels are elevated in the absence of CELF1

Among the 11 high-priority gene ligands of CELF1, we previously demonstrated that *Pax6* and *Prox1* are down-regulated by CELF1 in mouse lenses [35]. For further analysis, we focused on two additional genes with distinct molecular functions: *Gja8*, which encodes connexin 50, a transmembrane protein, and *Maf*, which encodes a nuclear transcription factor. In control postnatal day 15 (P15) lenses, GJA8 localizes along the membranes of lens fiber cells, which are organized in a regular pattern along the antero-posterior axis (Figure 5A, upper panels). In contrast, in P15 *Celf1*-cKO lenses, while GJA8 remains membrane-associated, the cellular architecture is disrupted. This disorganization is consistent with previously reported vacuolization and defective cell-to-cell junctions in *Celf1*-deficient lenses [32]. Moreover, GJA8 accumulates in distinct punctate structures rather than displaying its typical uniform distribution along the cell membranes. Notably, GJA8 signal intensity is markedly increased compared to controls, indicating that *Gja8* expression is higher in *Celf1*-cKO lenses (Figure 5A).

**Figure 5.**
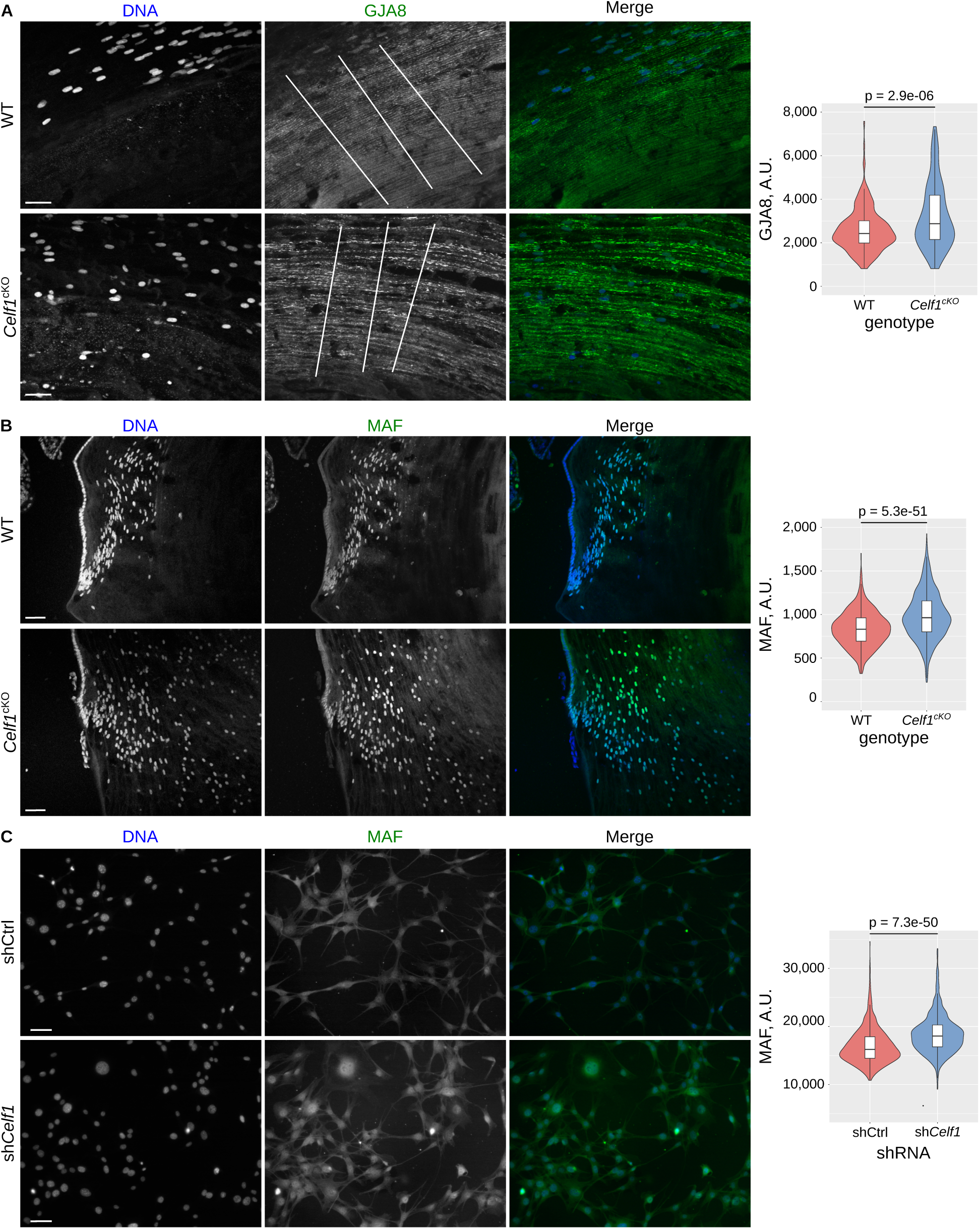
Expression of *Gja8* and *Maf* in mouse lenses and cultured lens cells **A**, Immunofluorescence of GJA8 in control (upper panel) and *Celf1*-cKO (lower panel) P15 mouse lenses. Right panel, quantification of the local maxima across the lines drawn in the second-left panel. **B**, Immunofluorescence of MAF in control (upper panel) and *Celf1*-cKO (lower panel) P15 mouse lenses. Right panel, quantification of MAF intensity in 300-500 DAPI-defined nuclei from several sections. **C**, Immunofluorescence of MAF in control (upper panel) and sh*Celf1* knockdown (lower panel) 21EM15 cells. Right panel, quantification of MAF signal intensity in 300-500 DAPI-defined nuclei from multiple fields. Scale bars 50 µm (**A**, **C**), 75 µm (**B**).

At the same developmental stage, MAF is absent from lens epithelial cells but begins to accumulate in the nuclei of cells entering the transition zone (Figure 5B). As cells move deeper into the lens and differentiate into fiber cells, MAF protein levels decline in maturing fiber cells. However, in *Celf1*-cKO lenses, MAF persists longer within internalized nuclei and displays stronger nuclear staining compared to controls, reflecting elevated MAF protein levels in the absence of CELF1 (Figure 5B).

To confirm these data, we used the 21EM15 mouse lens cell line, which includes a stable *Celf1* knock-down derivative [32]. As previously described [35], CELF1 depletion leads to increased PAX6 and PROX1 protein levels (Figure S2). While GJA8 was undetectable in these cells where the gene is not expressed, immunostaining confirmed elevated MAF protein levels in *Celf1*-depleted cells, paralleling the observations made in vivo (Figure 5C).

### Lens defects in *Xenopus* larvae overexpressing *maf*

Our data demonstrate that CELF1 directly binds to the *Maf* 3’UTR (Figure 1), that this 3’UTR represses translation in a CELF1-dependent manner (Figure 2), and that *Maf* is overexpressed in both cultured cells and lenses lacking CELF1 (Figure 5). To evaluate whether *Maf* overexpression contributes to the cataract phenotype observed in *Celf1*-cKO mice [32], we turned to *Xenopus laevis*, a well-established model for lens development and disease [55].

We injected *Xenopus laevis* embryos at the 4-cell stage with *Xenopus laevis maf* mRNA together with a lineage tracer. After two days of development, embryos were sorted based on tracer fluorescence to identify the injected side. Larvae were analyzed at 3 days post-fertilization. Figures 6A-C illustrate representative phenotypes. Overall body morphology appears normal following *maf* overexpression. The non-injected eyes are comparable to those of uninjected controls and tracer-only injected embryos (Figure 6D). In contrast, eyes on the injected side display various external defects, including microphtalmia with a greyish lens (Figure 6A) a ventral retinal gap suggestive of coloboma (Figure 6B), or a small, recessed eye (Figure 6C). Quantitative analysis of multiple larvae revealed that approximately 70% of *maf*-injected larvae exhibited externally visible eye abnormalities, compared to a low percentage in tracer-only controls (Figure 5E).

**Figure 6.**
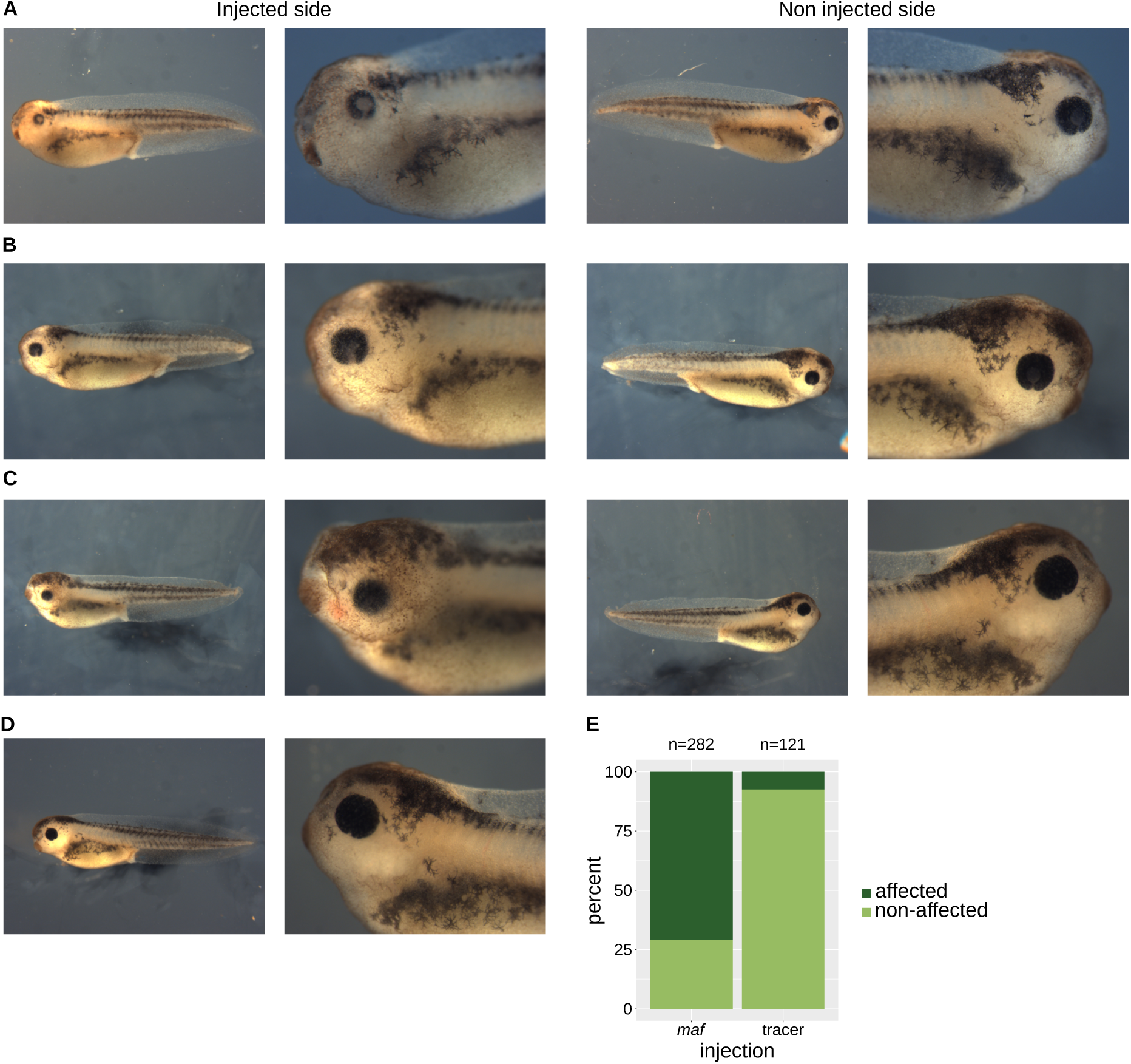
External morphology of *Xenopus laevis* larvae overexpressing *maf* **A-C**, External views of representative *Xenopus* larvae injected at the four-cell stage into one dorsal blastomere with mRNA encoding *Maf* and a fluorescent lineage tracer. Left panels, injected side (identified by fluorescence); right panels, uninjected control side. **D**, External view of a sibling embryo injected with lineage tracer only (injected side). **E**, Quantification of eye defects. p = 5.1e-31, chi-square test.

To further characterize the defects induced by *maf* overexpression, we performed histological analysis of larvae injected with lineage tracer alone or with *maf* mRNA (Figures 7A-D). In tracer only larvae, eye morphology and lens structure are similar on both sides, confirming that the lineage tracer does not interfere with lens development (Figure 7A). In contrast, while the non-injected sides of *maf*-injected embryos resemble those of controls (Figures 7B, C, D), the injected sides display notable abnormalities (Figures 7B’, C’, D’). In some cases, retinal layering is preserved (Figure 7B’), whereas in others it is severely disorganized (Figures 7C’, D’). Lens defects are consistently more severe, in that the injected sides exhibit smaller lenses (Figure 7B’), impaired fiber cell differentiation as evidenced by reduced eosin staining (Figure 7C’), or nearly complete lens absence, with only rudimentary remains (Figure 7D’). To quantify these defects, we measured eye and lens areas on serial sections by calculating the product of the minor and major diameters. Both are significantly reduced on the injected side (Figures 7E-F); however, lens growth is more severely impaired than overall eye growth, as reflected by a significantly lower lens-to-eye area ratio (Figure 7G).

**Figure 7.**
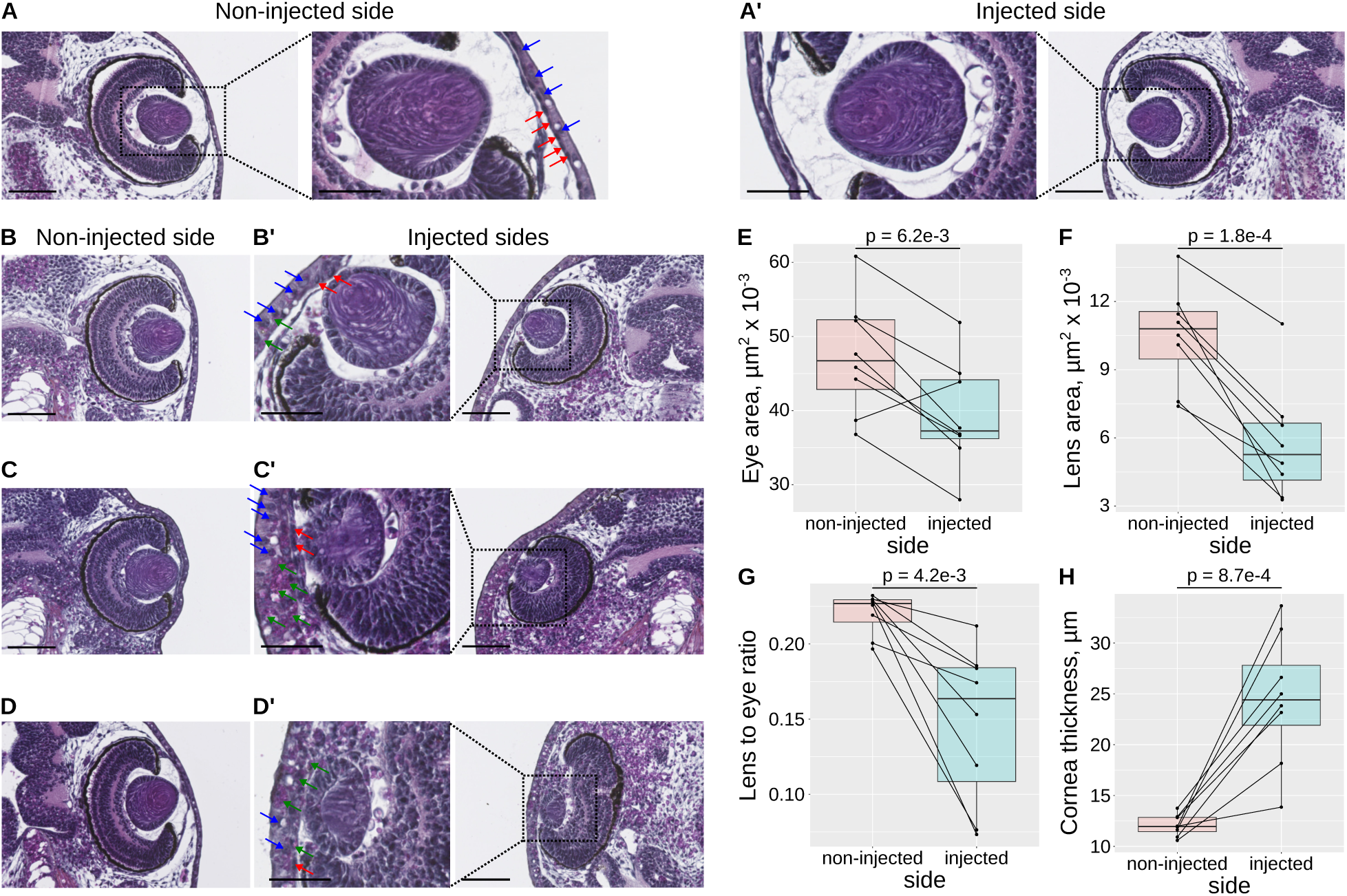
*maf* overexpression disrupts lens and corneal structure in *Xenopus laevis* larvae A-A’, Histological sections of a *Xenopus* larva injected at the four-cell stage into one dorsal blastomere with the lineage tracer alone. **A**, Non-injected control side. **A’**, Injected side. **B-D’**, Representative sections of sibling larvae co-injected with *maf* mRNA and the lineage tracer. In **B’, C’, D’**, higher magnification, arrow indicate: blue, nuclei of squamous corneal epithelial cells; red, nuclei of corneal presumptive endothelial cells; green, infiltrated cells. Scale bars 100 µm (low magnification), 50 µm (high magnification). **E-H**, quantifications of eye area (**E**), lens area (**F**), lens-to-eye area ratio (**G**), and corneal thickness (**H**), on injected vs non-injected sides (N = 8 larvae). p, paired Student’s t-test.

In addition to reduced eye and lens size, *maf* overexpression disrupts the organization of the presumptive cornea, the tissues separating the lens from the external environment. In tracer-only controls, the cornea displays a clearly defined outer squamous epithelium (blue arrows) and an inner presumptive endothelium (red arrows, Figure 7A), consistent with previous reports [56]. Figures 7B’, 7C’ and 7D’ show that this organization is lost in *maf*-injected larvae, which exhibit a prismatic corneal epithelium, a fragmented or discontinuous endothelium, and dense cellular infiltration (green arrows). We also observed consistent thickening of the presumptive cornea (Figure 7H), likely contributing to the externally visible “sunken” eye phenotype (e.g., Figure 6C). Previous studies using surgical or toxin-induced lens ablation have shown that lens-derived signals are essential for corneal differentiation [57–59]. Several pathways, including Semaphorin and TGF-β signaling, have been implicated [60,61]. Notably, lens-specific overexpression of TGF-β1 induces corneal thickening [61]. Therefore, the corneal defects seen in *maf*-overexpressing Xenopus larvae likely result from disrupted lens development. Collectively, these findings demonstrate that *maf* overexpression disrupts lens development in *Xenopus*, supporting the hypothesis that *Maf* derepression in *Celf1*-deficient mouse lenses contributes to cataract formation.

## DISCUSSION

Several human genetic studies have established a strong association between mutations in the *MAF* gene and congenital cataracts [62–72]. These mutations are typically point mutations located within the MAF DNA-binding domain and are often inherited in an autosomal dominant manner. In addition to cataracts, affected individuals frequently exhibit broader ocular developmental abnormalities, including Peters anomaly, Aymé-Gripp Syndrome, coloboma, amblyopa, or microcornea. In mouse models, two independent *Maf* mutations causing congenital cataracts were also found to be dominant or semi-dominant [73,74]. This supports the notion that cataract-associated *MAF* mutations are predominantly gain-of-function. This is exemplified by the D90V mutation in mice, which results in increased promoter activity of target genes [73].

Beyond these naturally occuring mutations, targeted gene inactivation strategies have been employed in mice to elucidate the role of *Maf* in lens development. Germline *Maf* knockout mice exhibit microphtalmia, with severe defects in lens morphogenesis resulting from impaired fiber cell elongation, and reduced expression of crystallin genes [75,76]. This underscores MAF’s function as a transcriptional regulator of crystallin genes, a conclusion further supported by luciferase reporter assays demonstrating that wild-type MAF, but not disease-associated mutants, activates promoters of four crystallin genes *CRYAA*, *CRYBA4*, *CRYBA1* and *CRYGA* [65]. Conditional inactivation of *Maf* in adult mice using a tamoxifen-inducible Cre system also leads to cataract formation, although with reduced severity compared to germline deletion [77]. This indicates that MAF is critical not only during lens development but also for maintaining lens structures in adults. Hence, both gain-of-function and loss-of-function (targeted disruption) mutations of *MAF* cause lens diseases in mammals, including cataracts.

Here, using *Xenopus* as a model system, we show that overexpression or ectopic expression of wild-type *maf* via mRNA injection is sufficient to induce lens defects. While gene overexpression may appear mechanistically similar to gain-of-function mutations, it can also have inhibitory effects by disturbing dosage-sensitive gene networks or competing for cofactors [78]. Thus, our finding that overexpression of wild-type *maf* leads to lens malformations unveils a distinct pathogenic mechanism involving *MAF* in cataract, complementary to traditional loss-of-function or point mutation models.

This raises an important question: to what extent can gene overexpression or ectopic expression be generalized as a pathogenic mechanism beyond *Maf*? We propose that the systematic identification of CELF1-bound RNAs in the lens, as reported in this study, provides a valuable set of candidates to address this question. We demonstrate for the first time, at a transcriptome-wide level and in a specific organ, that CELF1 functions as a global post-transcriptional repressor, reducing both mRNA abundance and translation efficiency. As a result, CELF1 deficiency leads to widespread upregulation of its mRNA ligands, including *Maf*. It is reasonable to hypothesize that overexpression of at least a subset of these ligands, especially those with known roles in lens development, may contribute with *Maf* to cataract formation in *Celf1* conditional knockout mice. A mouse model for ectopic expression of *Foxe3* (driven by the *Cryaa* promoter) in lens fiber cells has been analyzed by microarrays to identify the impact of its overexpression on the transcriptome [79]. A similar model for *Maf* would inform on the impact of its overexpression on the lens. Further, such a model would also provide new insights into the mRNAs misexpressed in *Celf1*^cKO^ lens, by indicating the mRNA targets that can be attributed to CELF1-based direct control, and the mRNA targets that likely result from indirect control by CELF1, via overexpression of *Maf*. Future studies that systematically test the impact of overexpressing individual CELF1 targets in model systems like *Xenopus* or lens organoids [55,80] could uncover novel genes whose dosage must be tightly regulated to maintain lens integrity. Such efforts would expand our understanding of lens biology and cataract pathogenesis, moving beyond conventional gene disruption studies and providing a new framework for functional validation of cataract candidate genes.

## MATERIALS AND METHODS

### Ethics statement

Mouse experiments were conducted either at the University of Rennes (France) or the University of Delaware (USA). In Rennes, mice were bred in the Biosit animal facility (Arche) approved by the French animal care agency (Direction des Services Vétérinaires, approval number A3523840). All procedures were performed following standard guidelines. At the University of Delaware, mice were bred and maintened in the Center for Animal Research facility. All animal protocols were reviewed and approved by the Institutional Animal Care and Use Committee (IACUC, protocol number 1226). All procedures conformed to the ARVO Statement for the Use of Animals in Ophthalmic and Vision Research.

*Xenopus laevis* experiments were conducted at the University of Rennes. Animals were housed in the IGDR facilities, accredited by the French animal care agency (Direction des Services Vétérinaires). Procedures were approved by the local ethics committee and authorized by the Ministry of Research (APAFIS #45521, 2023).

### iCLIP of CELF1

Lenses were dissected from euthanized 2-month-old mice, placed on Parafilm on ice, UV-irradiated (254 nm, 3 × 4000 µJ/cm²), ground in 100 µl PXL buffer per lens (50mM Tris-HCL pH 7.4, 0.1M NaCl, 1%NP40, 0.1% SDS, 0.5% DOC, plus DTT, RNasin and protease inhibitors Sigma P8340) and stored at −80°C. Dissections and irradiations were performed on 25 mice per iCLIP experiment, one mouse at a time to preserve lens freshness. The subsequent steps were essentially as described [37], except that the L3 primer was directly radiolabeled and adenylated to give L3-App via (1) phosphorylation with [γ-^32^P] ATP and PNK (New England Biolabs), and (2) adenylation of radiolabelled L3 with ATP and Mth RNA ligase (New England Biolabs). Briefly, lenses lysates were treated with Turbo DNase and RNase T1 (CLIP1: 2.4 U RNase T1 / mg protein extract, Sigma; CLIP2: equivalent of 100 U RNase T1 / mg protein extract, Ambion) for 10 minutes at 37°C. CELF1-RNA complexes were immunoprecipitated at 4°C with the 3B1 monoclonal antibody (Santa Cruz sc-20003), dephosphorylated, and ligated to radiolabeled, pre-adenylated L3-App. The 5′ end labeling step was omitted. Next, RNA-protein complexes were resolved on NuPAGE Bis-Tris gels, transfered to pure nitrocellulose membranes, and RNA was recovered from the membrane by proteinase K digestion. Reverse transcription, circularization-linearization, and PCR amplification were performed as described [37]. Libraries were sequenced on an Illumina platform (strand-specific, single-end, 50 bp reads).

### Analysis of iCLIP data and statistical analyses

Raw reads were processed using iCount 2.0.1 (https://icount.dev/). Reads were trimmed to remove adaptors, demultiplexed and filtered to retain those ≥ 19 nucleotides. Mapping was performed against the mouse genome (GRCm38) using STAR [81], keeping only uniquely mapped reads. Cross-link sites were defined as the terminal positions of the reads and scored by read count. Significance was calculated as in FOX2 studies [39], using a false discovery rate (FDR) calculated over ±3 nt windows compared to random background. Only sites with FDR < 0.05 were retained and termed *peaks*. Peaks <20 nt apart were clustered, and cluster scores were calculated as the sum of peak and adjacent signal scores. The data have been deposited in NCBI’s Gene Expression Omnibus and are accessible as GSE313211.

CLIP-seq data, as well as other quantitative data, were analyzed with R. Plots were prepared with the ggplot2 package. A fully annotated script is available upon request to the authors.

### CompBio-based analysis of iCLIP targets

A computational tool CompBio (developed at Washington University (GTAC@MGI, WashU School of Medicine, https://gtac-compbio-ex.wustl.edu—Academic/Non-Profit; PercayAI Inc, www.percayai.com/compbio-Commercial) was used to apply language processing and a curated biological dictionary to extract the concepts enriched in the candidates identified by the iCLIP assays [43–47]. CompBio performs de novo anlaysis of a given dataset using publicly available scientific literature (currently covering >30 million PubMed abstracts and >3 million full-text articles). CompBio reveals the biological “concepts” that are enriched in the dataset. It also identifies overarching “themes” clusters co-occurring concepts. As previously described, a normalized enrichment score (NEScore) indicates the significance of the identified themes and statistical confidence in the enrichment of a theme relative to random gene sets is indicated by P values. CompBio was used to analyze 689 genes (293 commonly found in both CLIP1 and CLIP2 plus those exclusively found in CLIP1 (295) and CLIP2 (101)). Themes that had NEScore ≥ 1.3 and P < 0.1 were considered significant. An automated theme annotation feature in CompBio was used (and modified where necessary) to underscore key biological processes.

### Luciferase assays

3′UTRs were PCR-amplified from reverse-transcribed mRNA of mouse lenses using gene-specific primers (Table S2). PCR products were cloned downstream of the firefly luciferase gene in the *pmirGlo* vector (Promega), using Gibson Assembly (NEBE2611S/L). Mutant constructs lacking CELF1-binding clusters were generated by amplifying 5’ and 3’ fragments separately (primer sequences in Table S2). Both fragments were inserted simultaneously into *pmirGlo*.

Transfections and luciferase assays followed previously described methods [30].

### Immunofluorescence on mouse eye

Eyes from postnatal day 15 (P15) were fixed in 4% paraformaldehyde (PFA) for 30 min on ice, followed by incubation in 30% sucrose overnight at 4°C, embedded in OCT (Tissue-Tech), and sectioned at 14 μm. Sections were blocked in 5% chicken serum (Abcam, Cambridge, UK) with 0.3% Triton for 1 hour at room temperature. The sections were incubated overnight at 4°C with the following primary antibodies: CELF1 (Santa Cruz Sc-20003, 1:500), MAF (Abclonal A12720, 1:2500), GJA8 (Life Technologies PA5-11644, 1:1000). Sections were washed and incubated with Alexa Fluor 594-conjugated secondary antibody (1:1000; Life Technologies, Carlsbad, CA) and DAPI (1:1000) for 1 hr at room temperature. Slides were mounted using mounting media (ProLong Gold antifade reagent, Invitrogen P36930) and imaged on a Zeiss LSM 880 confocal microscope configured with Diode/Argon laser (405 nm, 488 nm and 594 excitation lines). Image brightness/contrast was adjusted identically across all conditions using Adobe Photoshop. Signal quantification was performed using Fiji (ImageJ). Data represent three biological replicates per condition.

### Cell culture and immunocytofluorescence

21EM15 mouse lense epithelial cells (from Dr. John Reddan, Oakland University, Rochester, MI, USA) were cultured in DMEM with 4.5 g/L glucose, L-glutamine and sodium pyruvate (Life Technologies, Carlsbad, CA, USA) with 10% FBS (Eurobio, Les Ulis, France) and 1% penicillin-streptomycin (Life Technologies, Carlsbad, CA, USA) at 37 °C, 5% CO2. Cells were split three times weekly and typically reached 80% confluence within three days. The stable *Celf1*-knockdown 21EM15 derivative, generated via lentiviral transduction with *Celf1*-specific shRNA, has been described [32].

For immunostaining, 5×10^4^ cells/cm^2^cells were seeded on glass coverslips. After 24 hours, cells were fixed in 4% PFA in MES buffer (10 mM MES buffer, 138 mM KCl, 3 mM MgCl _2_, 0.1 g/ml sucrose and 2 mM EGTA, pH 6.1) for 10 minutes. They were permeabilized with 0.1% Triton X-100, and blocked in 1% BSA in PBST for 1 h at room temperature. Primary antibodies in PBS-BSA include Rabbit anti-MAF (1:500; Abcam 77071), rabbit anti-PAX6 (1:200; Abcam Ab2237), rabbit anti-PROX1 (1:200; Biolegend, 925201). The cells were incubated with antibodies overnight at 4°C. After washes, cells were incubated with the secondary antibodies (Alexa Fluor 488 or 555 anti-rabbit; 1:1000, Invitrogen) for 1 h at RT. Coverslips were mounted with ProLong Gold (Invitrogen) containing 1 µg/mL DAPI (Sigma-Aldrich). Imaging was done using a Nikon NIE microscope with a 40× objective.

### Xenopus embryo manipulations and histological methods

*Xenopus laevis maf.L* ORF was PCR-amplified from reverse-transcribed Xenopus embryo mRNA with the primers given in Table S2. The PCR product was cloned into BglII-NotI cleaved pT7TS vector [29], using Gibson Assembly (NEBE2611S/L). The plasmid was linearized with BamH1 before in vitro transcription (Message Machine T7, Life Technologies AM1344). One dorsal blastomere of four-cell *Xenopus laevis* embryos was injected with 1.25 ng Alexa 568-conjugated dextran (Life Technologies D22912) and 1.5 fmol *maf* mRNA. Embryos were sorted according to injection side after two days at 22°C, and analysed at day 3 for eye malformation.

For histology, embryos at stages 39/40 were fixed in Carnoy for 2 hours at room temperature. They were dehydrated (Ethanol 100% Eosin 20 minutes, Ethanol 100% 20 minutes twice, Butanol 30 minutes twice) and paraffin-embedded as described [55]. Horizontal sections (7 µm) were stained with Hematoxylin and Eosin.

## Supporting information

Legends to the supplemental tables

Supplemental figures with legends

## ACKNOWLEDGMENTS

We thank Vincent Legagneux for his help during the preparation of biological samples for iCLIP-seq. We thank the Biosit platforms (Rennes, France) for giving us access to their facilities, and notably Arche (mouse husbandry), MRic (microscopy) and H2P2 (histology).

This work was supported by a grant from the Agence Nationale de la Recherche [ANR SILENS, ANR-24-CE14-3347] to LP, and by grants from the National Institutes of Health [R01 EY021505 and R01 EY036923] to SAL.

