## Supplementary material for "Mapping of CELF1-RNA interactions reveals post-transcriptional control of lens development": Legends to the supplemental tables

### LEGENDS TO SUPPLEMENTARY TABLES

**Table S1. iCLIP-seq data**

Total score, sum of all cluster scores within the 3'UTR. maxUGU, number of UGU within the 3'UTR cluster containing the highest number of UGU. Exp, expression (CPM). Pubmed, number of citations combining gene name with the keyword "lens". HeLa, gene previously identified as a CELF1 ligand in HeLa cells [41]. catmap, gene present in the Cat-Map database, February 2025 release [10,11]. isyte, maximal enrichment score in iSyTE 2.0 [50].

**Table S2. Primer sequences**

Sequences of the primers used for constructs, with their usage.
