## Supplemental figures with legends for "Mapping of CELF1-RNA interactions reveals post-transcriptional control of lens development"

### FIGURE S1

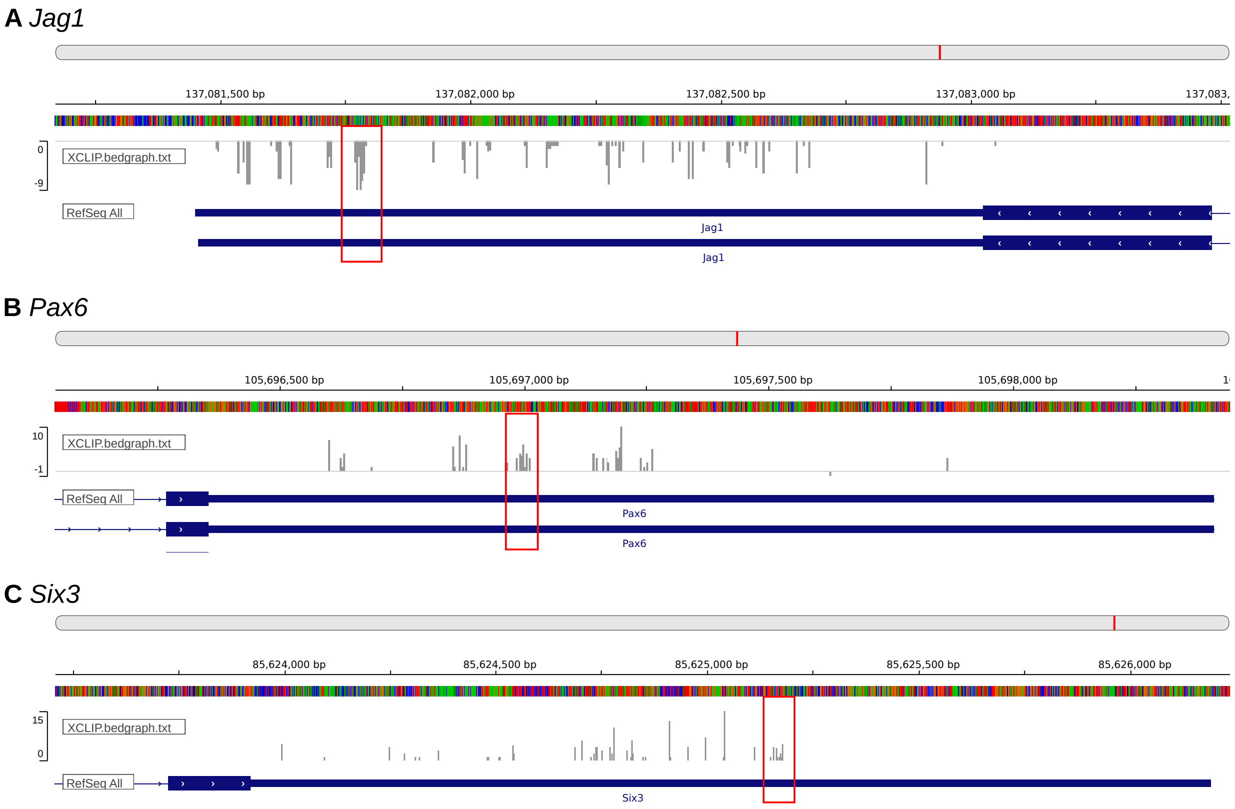


**Figure S1. CELF1 binding clusters in *Jag1*, *Pax6*, *Six3***

CELF1 binding clusters identified in the 3'UTRs of *Jag1* (**A**), *Pax6* (**B**) and *Six3* (**C**). Red rectangles indicated high-density regions of CELF1-binding clusters that were deleted in the mutant constructs used in Figure 2E.

#### FIGURE S2

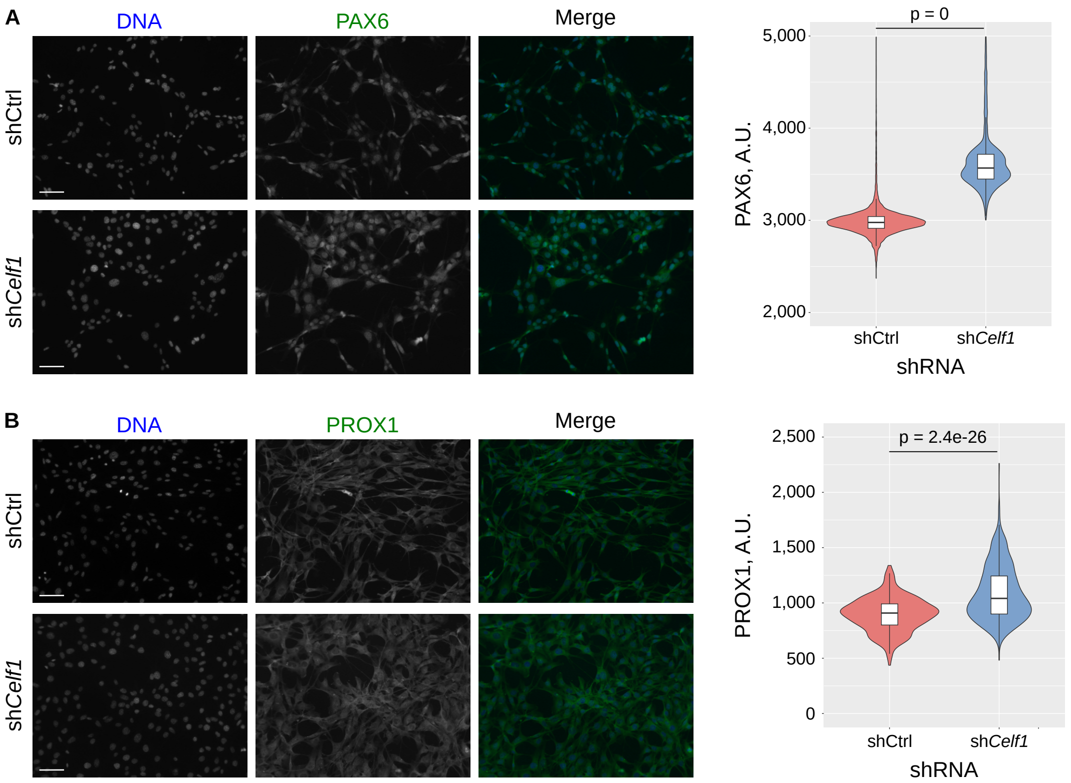


**Figure S2. Expression of *Gja8* and *Maf* in mouse lenses and cultured lens cells**
Immunofluorescence of PAX6 (**A**) and PROX1 (**B**) in control (upper panel) and sh*Celf1* knockdown (lower panel) 21EM15 cells. Right panel, quantification of PAX6 and PROX1 intensities in 300-500 DAPI-defined nuclei across multiple fields. Scale bars 50 µm.
